# Two common disease-associated TYK2 variants impact exon splicing and TYK2 dosage

**DOI:** 10.1101/830232

**Authors:** Zhi Li, Maxime Rotival, Etienne Patin, Frédérique Michel, Sandra Pellegrini

## Abstract

TYK2 belongs to the JAK protein tyrosine kinase family and mediates signaling of numerous antiviral and immunoregulatory cytokines (type I and type III IFNs, IL-10, IL-12, IL-22, IL-23) in immune and non-immune cells. After many years of genetic association studies, *TYK2* is recognized as a susceptibility gene for some inflammatory and autoimmune diseases (AID). Seven *TYK2* variants have been associated with AIDs in Europeans, and establishing their causality remains challenging. Previous work showed that a protective variant (P1104A) is hypomorphic and also a risk allele for mycobacterial infection. Here, we have studied two AID-associated common TYK2 variants: rs12720270 located in intron 7 and rs2304256, a non-synonymous variant in exon 8 that causes a valine to phenylalanine substitution (c.1084 G > T, Val362Phe). We found that this amino acid substitution does not alter TYK2 expression, catalytic activity or ability to relay signaling in EBV-B cell lines or in reconstituted TYK2-null cells. Based on *in silico* predictions that these variants may impact splicing of exon 8, we: i) analyzed *TYK2* transcripts in genotyped EBV-B cells and in CRISPR/Cas9-edited cells, ii) measured splicing using minigene assays, and iii) performed eQTL (expression quantitative trait locus) analysis of *TYK2* transcripts in primary monocytes and whole blood cells. Our results reveal that the two variants promote the inclusion of exon 8, which, we demonstrate, is essential for TYK2 binding to cognate receptors. In addition and in line with GTEx (Genetic Tissue Expression) data, our eQTL results show that rs2304256 mildly enhances *TYK2* expression in whole blood. In all, these findings suggest that these *TYK2* variants are not neutral but instead have a potential impact in AID.

## Introduction

Early genetic association studies have assigned to the *TYK2* locus an impact on susceptibility to systemic lupus erythematosus (SLE) and other autoimmune diseases (AID). The identification has been replicated in a number of recent analyses, and *TYK2* is now recognized as a susceptibility gene in a variety of inflammatory and autoimmune diseases, including type I diabetes (T1D), psoriasis and multiple sclerosis (Table 1). These chronic disorders have a complex etiology where combinations of genetic and environmental factors eventually lead to loss of immunological tolerance, chronic immune activation, and damage to one organ or several tissues [1].

**Table 1.**
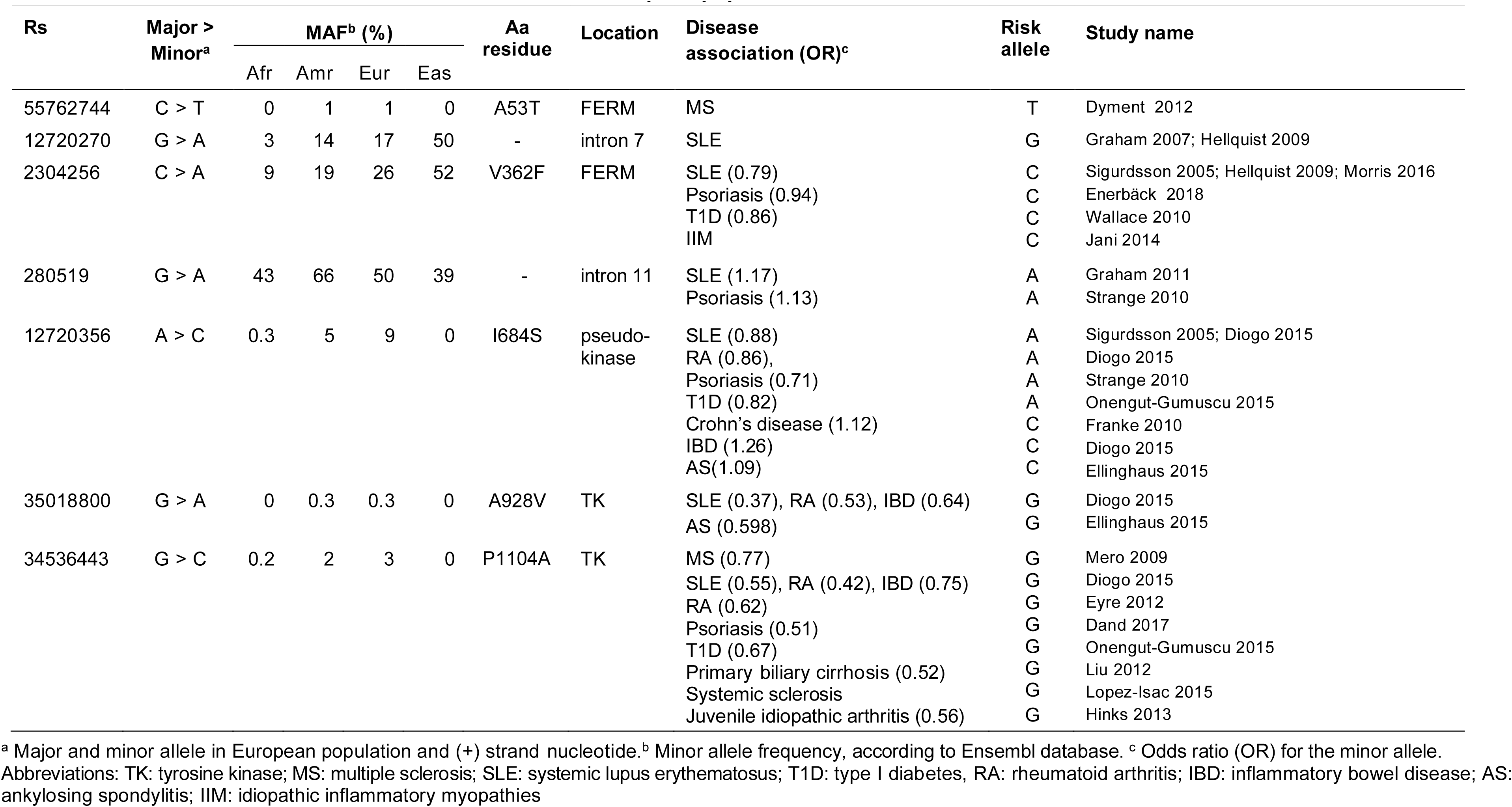
Autoimmune disease-associated TYK2 variants in European population.

How *TYK2* variants impact disease onset or progression remains an open question. In human populations, the *TYK2* locus presents with thousands of single nucleotide polymorphisms (SNP), of which more than 500 cause non-synonymous (amino acid-altering) changes. Seven *TYK2* variants have been associated with AIDs in European cohorts and for most the minor allele is protective (Table 1). Notably, rs12720356 (I684S) is protective for some AID but risky for others, which suggests different underlying pathogenic mechanism. Although these associations are of relatively weak magnitude, they may be relevant in view of promising development of small molecule selective TYK2 inhibitors to be used in the clinic [2]. Only the biological understanding of each association will enable patient stratification for individualized molecular targeted therapy. For this, it will be necessary to map causal SNP(s) within clustered SNPs that are in linkage disequilibrium (LD), validate their impact on TYK2 expression and function, define which signaling pathways and molecular processes are most affected by the variation and possibly identify the cell type(s) and/or cell state(s) driving the association in any given disease. These represent major challenges, since TYK2 is a ubiquitous tyrosine kinase that relays signaling of many antiviral and immunoregulatory cytokines (type I and type III IFNs, IL-10, IL-12, IL-22, IL-23) acting on a variety of immune and non-immune cells [3, 4].

Among the seven disease-associated *TYK2* SNPs, five cause a single amino acid change (Table 1). We have previously reported on rs12720356 and rs34536443, which map in the regulatory pseudo-kinase domain and the tyrosine kinase domain, respectively. Both protein variants, TYK2-I684S and TYK2-P1104A, are catalytically impaired but relay signaling in reconstituted non-immune cells [5]. Further studies in immune cells showed that rs34536443 (P1104A) homozygosity led to reduced type I IFN, IL-12 and IL-23 signaling [6], while rs12720356 (I684S) did not alter TYK2 function in cytokine signaling [5–7]. Boisson-Dupuis *et al* also found that homozygosity at rs34536443 confers predisposition to tuberculosis and strongly impairs IL-23 signaling in T cells and IFN-γ production in PBMC [7, 8]. Gorman *et al* reported that rs34536443 heterozygosity leads to reduced IFN-α signaling in naïve but not effector T cells, and that carriers have reduced circulating Tfh cells [9]. Another study showed that heterozygosity at rs12720356 (I684S), but not rs34536443 (P1104A), leads to reduced IL-12 signaling in CD4+ and CD8+ T cells [10]. While these findings underline a complex picture, they do converge on the view that these two hypomorphic alleles confer protection to auto-inflammatory and auto-immune conditions in European populations.

Here, we present a study of two additional disease-associated variants, which are in LD (r2=0.50 and 0.9 in European and Asian populations, respectively) (Ensembl/1000 Genomes). Rs12720270 has been associated to protection from SLE in UK families and in Finnish population [11, 12]. Rs2304256 has been associated to protection from SLE [12–14], psoriasis [10] and T1D [15] in Caucasian populations. Based on predictions that these polymorphisms may impact splicing of the flanking exon, we analyzed *TYK2* transcripts in EBV-B cells from genotyped donors. We also studied splicing using a conventional minigene assay and CRISPR/Cas9-edited cells. Combined to *cis* expression quantitative trait locus (eQTL) analysis of *TYK2* in monocytes and whole blood from genotyped individuals, our results revealed an impact of these variants on splicing of a small exon which is essential for TYK2 binding to cytokine receptors.

## Results

### Rs2304256 and rs12720270 promote exon 8 inclusion

Rs2304256 is a non-synonymous variant (C > A) that maps in *TYK2* exon 8 and causes a valine to phenylalanine substitution (c.1084 G > T, Val362Phe) (Fig 1). Valine 362 is located in the FERM domain, which, together with the SH2-like domain, mediates the interaction of TYK2 with cognate cytokine receptors, such as IFNAR1 and IL-12Rβ1. Through this interaction, TYK2 sustains the level of these receptors and calibrates cytokine signaling [16, 17]. Valine 362 is not evolutionarily conserved nor is found in the other members of the JAK family (Fig 1C) and, according to PolyPhen-2, substitution with a phenylalanine is predicted to be benign. To assess the functional consequence of this amino acid substitution, we measured IFN-α signaling in four EBV-B cell lines homozygous for the major allele (CC, TYK2-V362) and two lines homozygous for the minor allele (AA, TYK2-F362) (Fig 2A). All lines clearly responded to low level IFN-α, as seen by induced phosphorylation of TYK2 and STAT1, even though an inter-individual variability was observed. We thus turned to conventional studies of engineered TYK2 expressed in TYK2-null cells. TYK2-V362 and TYK2-F362 were transiently transfected in CRISPR/Cas9-edited TYK2-null 293T cells and IFN response was measured. As shown in Fig 2A, TYK2-V362 and TYK2-F362 were similarly expressed, rescued IFNAR1 level and relayed IFN-α-induced phosphorylation of TYK2 and STAT1 to the same extent (Fig 2B). Comparable catalytic activity was indicated by the basal P-TYK2 in non-treated samples (Fig 2B). To strengthen these findings, we studied TYK2-null fibrosarcoma 11,1 cells [18] transfected with TYK2-V362, TYK2-F362, variant TYK2-A928V, variant TYK2-P1104A (these latter in the F362 background) and the *bona fide* kinase-dead TYK2-K930R [19]. As shown in Fig 2C, TYK2-V362 and TYK2-F362 rescued IFN-α-induced signals. TYK2-K930R failed to rescue, while TYK2-A928V and TYK2-P1104A, protective in many autoimmune diseases, partially rescued signaling to IFN. Basal P-STAT1 reflects TYK2 catalytic activity in F362- or V362-expressing cells and was not detected in cells expressing catalytically impaired TYK2-A928V, TYK2-P1104A, TYK2-K930R. In sum, TYK2-V362 and TYK2-F362 did not exhibit functional difference. Thus, we considered the possibility that the nucleotide variation at rs2304256 affects pre-mRNA processing. It is well known that exonic sequences can impact gene expression and that natural variations or somatic mutations can modulate pre-mRNA processing [20]. In close proximity of the intron 7/exon 8 boundary, rs2304256 might not be neutral but may influence splicing efficiency of exon 8 (Fig 1A). An *in silico* analysis (Human Splicing Finder, http://www.umd.be/HSF) predicted that the C (major/ancestral) to A (minor/derived) nucleotide change at rs2304256 could destroy a putative exonic splicing enhancer (ESE) regulated by the SR protein SF2/ASF, generating a new ESE recognized by the SRp55 protein or breaking a silencer motif. In parallel, we also studied rs12720270, which is located in intron 7, 36 nt upstream of the intron 7/exon 8 boundary (Fig 1A). The *in silico* analysis of intron 7 sequence predicted that the G (major/ancestral) to A (minor/derived) nucleotide substitution could break a potential branch point and alter pre-mRNA splicing.

**Fig 1.**
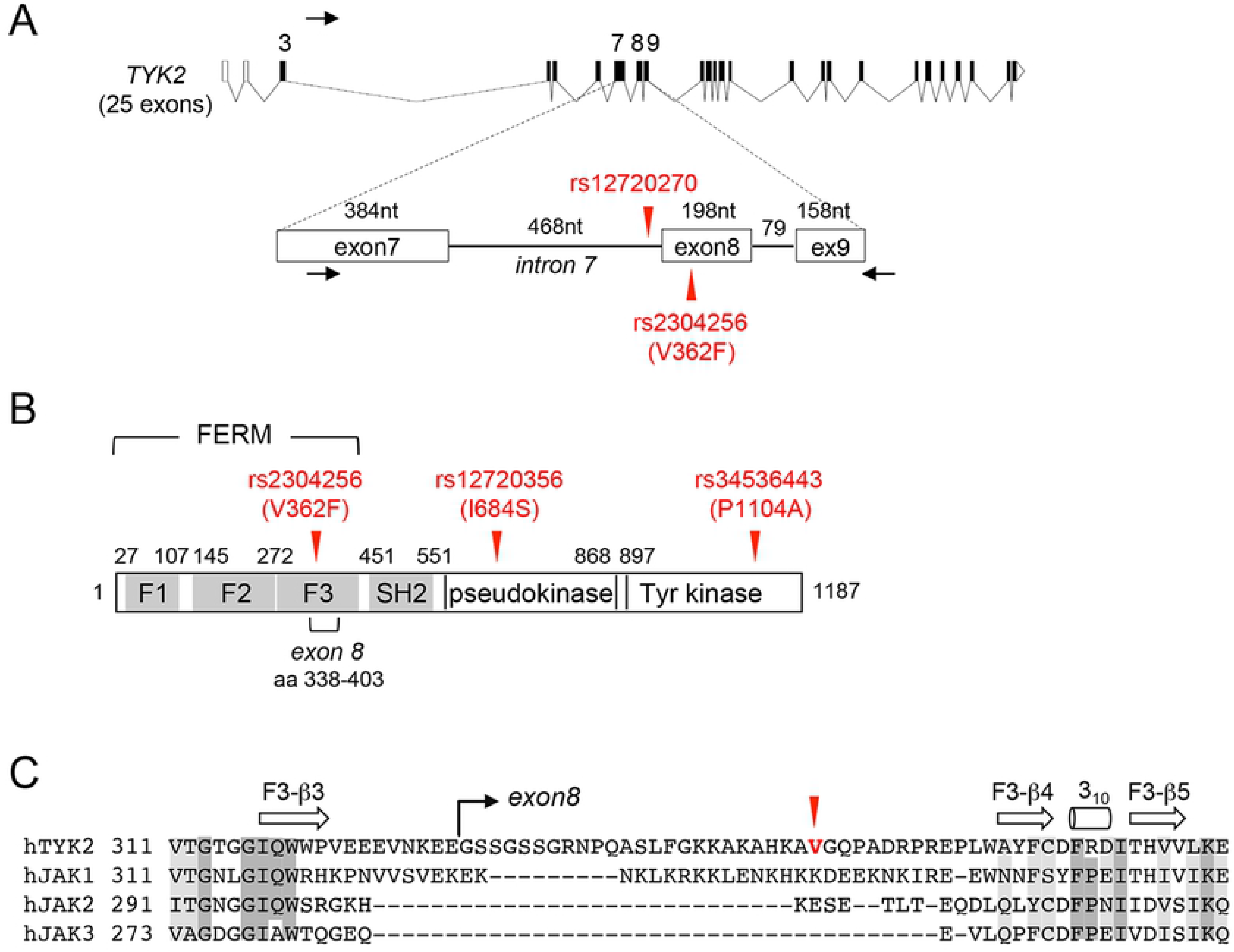
Position of rs12720270 and rs2304256 in *TYK2*. (A) The intron-exon organization of the *TYK2* locus is shown above. Coding exons are represented as filled boxes and introns as lines. Two 5’ non-coding exons are represented as empty boxes. The region from exon 7 to exon 9 is expanded below. Rs12720356 is located 36 nt upstream of the intron 7/exon 8 boundary. Rs2304256 is 75 nt downstream of the boundary. The black arrows point to the 5’ and 3’ primers used for PCR to amplify endogenous *TYK2* transcripts spanning exon 7 to exon 9. (B) Domain organization of the TYK2 protein. The FERM domain is made of 3 subdomains (F1 to F3). The position of V362F, I684S and P1104A encoded by rs2304256, rs12720356 and rs34536443, respectively, are arrowed in red. (C) Alignment of the four human JAK proteins in the region surrounding TYK2-Val362 in red. Secondary structures are indicated above, with β-strands as arrows and α-helices as cylinders.

**Fig 2.**
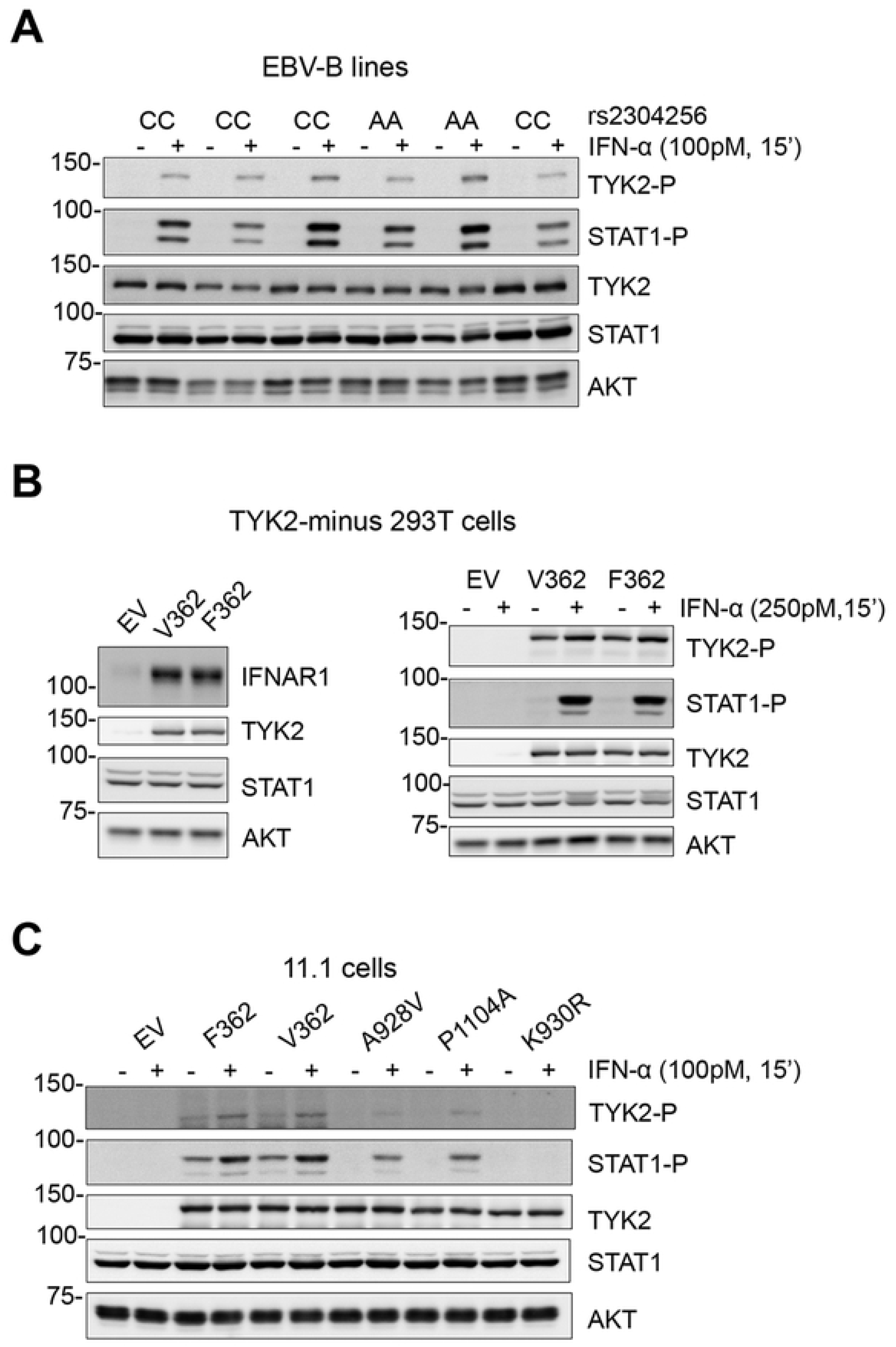
No impact of the V362F substitution on TYK2 function. (A) IFN-α-induced JAK/STAT activation in six EBV-B cell lines genotyped for rs2304256. Cells were treated with IFN-α for 15 min, TYK2 and STAT1 tyrosine phosphorylation was analysed by western blot with phospho-specific Abs. Membranes were reprobed for TYK2 and STAT1. (B) Level of IFNAR1 (left) and IFN-α-induced TYK2 and STAT1 phosphorylation (right) in TYK2-minus 293T cells transiently expressing TYK2-V362 or TYK2-F362 in the pIRES vector. EV, empty vector. (C) IFN-α-induced TYK2 and STAT1 phosphorylation in TYK2-deficient 11,1 cells transiently expressing TYK2-F362, TYK2-V362 or three other mutants (A928V, P1104A, K930R) in the F362 backbone. EV, empty pRc-CMV vector.

Based on the above, we asked whether rs12720270 and/or rs2304256 had any impact on transcripts at the level of exon 8 of *TYK2*. For this, we set up a RT-PCR assay with primers mapping in exon 7 and exon 9 of *TYK2* and used as template cDNA from 30 EBV-B cell lines (Table 2) (Materials and Methods). In addition to the expected product (740 nt), in some cell lines a minor product was amplified. Results for 15 representative cell lines is shown in Fig 3A. Sequencing showed that the minor product (542 nt) lacked 198 nt corresponding to exon 8 and retained the original open reading frame (Fig S1A). Interestingly, 12 of the 30 cell lines analyzed did not yield the 542 nt PCR product. Among these 12 lines, nine were homozygous for the minor rs2304256 allele (AA), one was homozygous for the minor rs12720270 allele (AA) and two were homozygous for both minor alleles (Table 2). These data suggested that minor alleles (homozygosity) at rs12720270 and/or rs2304256 promote the inclusion of exon 8. Analysis of PBMC samples purified from two genotyped donors confirmed this observation (Fig 3A).

**Table 2.** Analysis of exon 8 splicing in genotyped EBV-B cell lines.

| EBV-B<br>line | cell 550nt band<br>(Δexon 8) | Rs2304256<br>(V362F) | rs12720270<br>(intron 7) |
| --- | --- | --- | --- |
| 50044 | + | CC | GG |
| GM07056 | + | CC | GG |
| GM10854 | + | CC | GG |
| GM10846 | + | CC | GG |
| GM10857 | + | CC | GG |
| GM10851 | + | CC | GG |
| GM11830 | + | CC | GG |
| GM12004 | + | CC | GG |
| GM12282 | + | CC | GG |
| GM10836 | + | CA | GG |
| GM10859 | + | CA | GG |
| GM10864 | + | CA | GG |
| 51814 | + | CA | GG |
| GM12145 | + | CA | GG |
| GM11918 | + | CA | GA |
| GM10835 | + | CA | GA |
| GM12144 | + | CA | GA |
| 52173 | + | CA | GA |
| GM12249 | — | CA | AA |
| 34702 | — | AA | GG |
| GM12275 | — | AA | GG |
| 50772 | — | AA | GG |
| 51464 | — | AA | GG |
| GM11882 | — | AA | GG |
| 52279 | — | AA | GA |
| 49888 | — | AA | GA |
| GM07037 | — | AA | GA |
| GM12154 | — | AA | GA |
| GM12342 | — | AA | AA |
| GM12234 | — | AA | AA |

**Fig 3.**
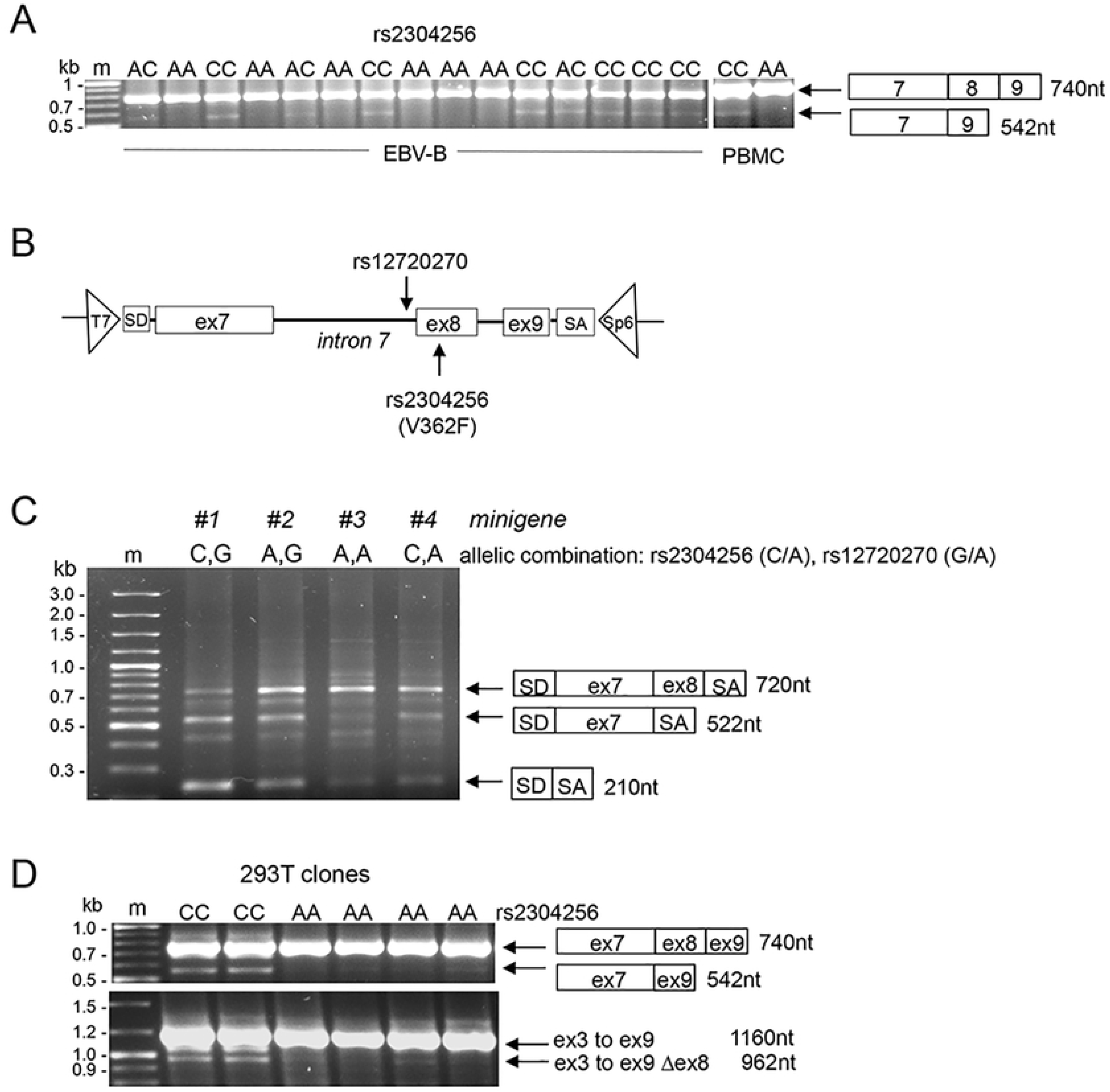
Analysis of the effect of the two variants on exon 8 splicing. (A) Analysis of *TYK2* exon 8 in 15 EBV-B lines and PBMC from individuals genotyped at rs2304256. RT-PCR assay was performed with primers mapping in *TYK2* exon 7 and exon 9 (see Fig 1A). See also Table 2. (B) Schematics of *TYK2* genomic sequence from exon 7 to exon 9 inserted into the pI-12 vector used for minigene analysis. The positions of the two variations is indicated. The vector contains splice donor site (SD) and splice acceptor site (SA). Minigene-specific primers (T7 and Sp6) used for PCR and sequencing are indicated by the triangles. (C) A representative result of the analysis of the four minigenes in 293 cells. Exon 9 is not expressed probably due to the weak acceptor site at its 5’ and the strong SA site of the vector. (D) Analysis of *TYK2* exon 8 in 293T clones edited at rs2304256 by CRISPR/Cas9. Two unedited clones (CC) and four edited clones (AA) were analyzed. Top panel, RT-PCR was performed as described in (A). Bottom panel, RT-PCR using a forward primer in exon 3 and a reverse primer in exon 9 (Fig 1A). The middle band is a product resulting from aberrant pairing of the two other products.

To further assess the effect of the nucleotide variations on exon 8 and exclude potential influence of other polymorphisms on transcript abundance or splicing, we turned to the minigene assay (Cooper, Methods 2005). Using the pI-12 splicing vector, we generated a minigene reporter construct covering 1.2 kb of *TYK2* genomic DNA, from exon 7 to exon 9, and comprising rs12720270 and rs2304256 (Fig 3B, top). The four allelic combinations were generated and transfected in 293T cells. After RNA extraction and cDNAs synthesis, splicing products were analysed by PCR using T7 and Sp6 primers specific to the minigene (Fig 3B, bottom). The three major amplification products obtained were sequenced. The 720 nt product contained exon 7 and exon 8. Of note, exon 7 of the minigene was about 70 nt shorter than the *bona fide* exon 7 due to the use of an alternative acceptor site. Also, the absence of exon 9 most likely reflects the weakness of its acceptor site with respect to the strong splicing acceptor site (SA) present in the vector. The 520 nt product contained only exon 7. The 210 nt product contained the 5’ exon (SD) and 3’ exon (SA) of the pI-12 vector. In comparing the profile of transcripts expressed from the four minigenes, it appeared that, for each of the two SNPs, the presence of the minor allele A led to a higher level of a transcript retaining exon 8 (Fig 3B). Moreover, the minigene #3 carrying both minor alleles A almost exclusively expressed the exon8-inclusive transcript.

293T cells are homozygous for the rs2304256 major allele. Using the CRISPR/Cas9 genome editing technology, we obtained clones homozygous for the minor A allele. RNAs from four edited clones (AA) were analyzed by RT-PCR as described above. The PCR product lacking exon 8 (550 nt) was undetectable in three edited clones and barely detectable in the fourth clone (Fig 3C, top). These results were corroborated by using a set of primers covering from exon 3 (the first coding exon) to exon 9. Sequencing of the products confirmed that the smaller transcript (960 nt) amplified from control cells lacked exon 8 (Fig 3C, bottom panel).

### TYK2-ΔE8 is catalytically competent but unable to mediate cytokine signaling

The results described above indicated that rs2304256 and rs12720270 influence pre-mRNA processing at the level of exon 8. Interestingly, one annotated *TYK2* transcript (*TYK2-204*) lacks exon 8 (ENST00000525220). Both *TYK2-204* and the small transcript detected in EBV-B cells retain the correct ORF and may give rise to a protein isoform missing the exon 8-encoded segment. We did not succeed in detecting such protein probably owing to its low expression level. Yet, the alternate transcript - and protein - may be more abundantly expressed in specific cell types and/or under particular conditions. On this basis, we undertook functional studies of a TYK2-ΔE8 protein. Our reference *TYK2* cDNA was deleted of the exon 8 sequence (198 nt, encoding aa 338-403). TYK2-WT and TYK2-ΔE8 expression vectors were stably transfected in TYK2-null cells and clones expressing similar TYK2 levels were chosen. We first compared the catalytic activity of WT and ΔE8. TYK2 was immunoprecipitated and subjected to *in vitro* kinase assay. As shown in Fig 4A, both proteins were basally phosphorylated in cells (lanes 1 and 3) and, when ATP was added to the reaction, the intensity of the phospho-TYK2-reactive bands increased (lanes 2 and 4), demonstrating that TYK2-ΔE8 is as catalytically competent as the wild type protein. Next, we compared the ability of TYK2-ΔE8 to rescue cytokine signaling. The level of TYK2 and STATs phosphorylation was measured in cells pulsed with IFN-β. Fig 4B shows that two TYK2-ΔE8-expressing clones were totally unresponsive to cytokine stimulation, as compared to two WT-expressing clones. Since a key function of TYK2 is to sustain cognate receptors, i.e. IFNAR1, IL-12Rβ1 and IL-10R2 [17, 21], we measured IFNAR1 levels in TYK2-null cells (lane 1) and in the derived clones. Fig 4C shows that TYK2-ΔE8 is unable to exert its scaffolding function and rescue IFNAR1 levels.

**Fig 4.**
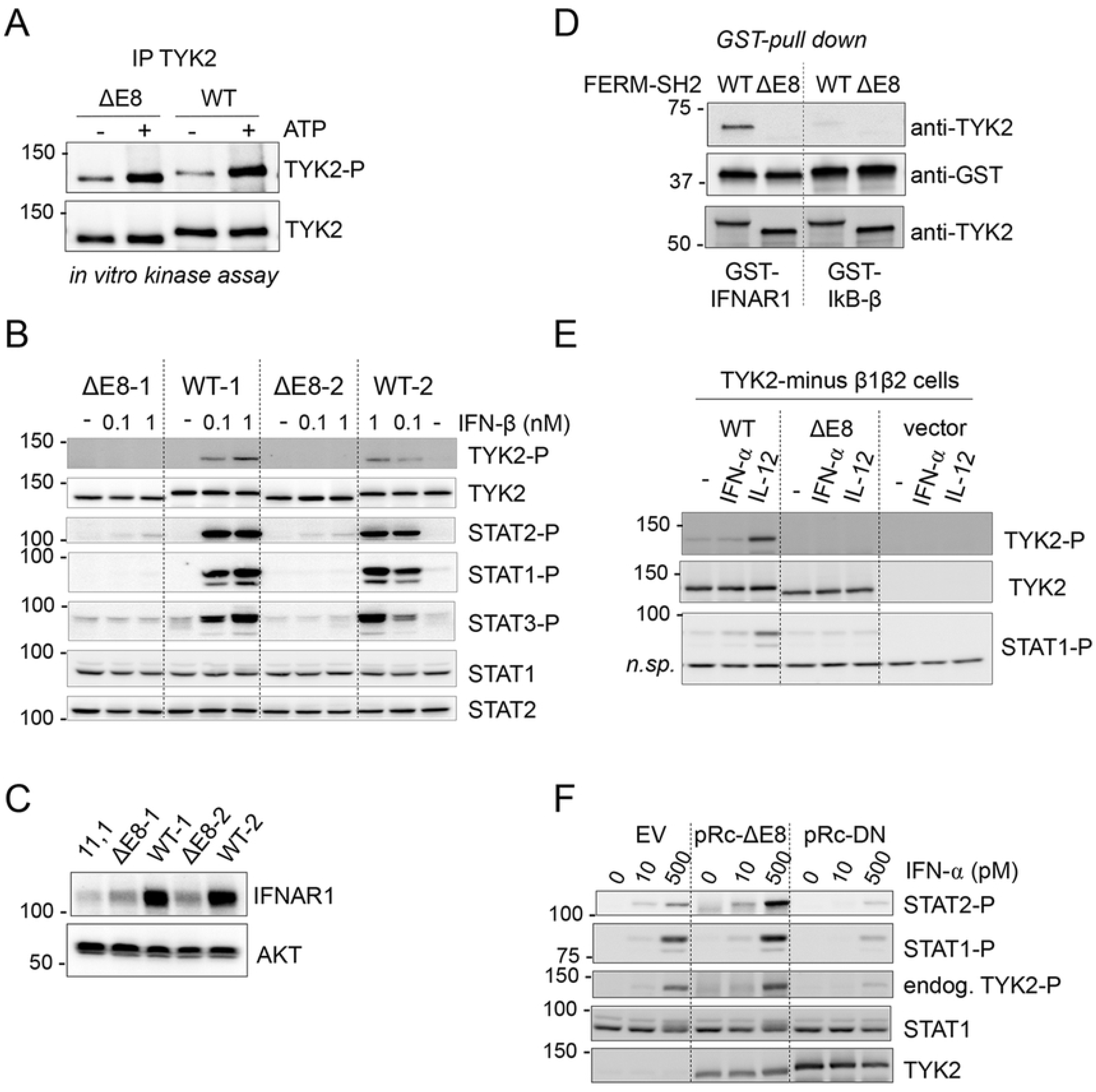
TYK2-ΔE8 is catalytically active but unable to rescue cytokine signaling. (A) Basal *in vitro* kinase activity of TYK2 from unstimulated 11,1 cells stably expressing TYK2 WT or TYK2-ΔE8. TYK2 was immunoprecipitated and subjected to an *in vitro* kinase reaction for 5 min at 30°C in the presence (+) or absence (-) of 30 μM ATP. Phosphorylated TYK2 in the reaction was revealed by immunoblotting with anti-phospho-TYK2. The membrane was reprobed for TYK2. (B) IFN-induced JAK/STAT activation in 11,1 cells stably expressing TYK2 WT or TYK2-ΔE8. Cells were treated with IFN-β for 15 min. The level of tyrosine-phosphorylated TYK2, STAT1, STAT2 and STAT3 was analyzed by with phospho-specific Abs. The membrane was reprobed for TYK2 and total STATs. (C) IFNAR1 level in 11,1 cells (-) and derived clones stably expressing TYK2 WT or TYK2-ΔE8. (D) *In vitro* interaction of His-TYK2-FERM-SH2 with GST-IFNAR1cyt. His-TYK2-FERM-SH2 WT or ΔE8 were incubated with a GST fusion protein containing the cytoplasmic domain of IFNAR1(IFNAR1*cyt*) or IkB-β. Proteins bound to glutathione-Sepharose beads were separated on SDS-PAGE and visualized with TYK2 Abs. Five % input TYK2 protein shown at the bottom. (E) IL-12-induced JAK/STAT activation in 11,1 cells stably expressing the IL-12 receptor β1 and β2 chains. Cells were transiently transfected with TYK2 WT or TYK2-ΔE8. Twenty-four hrs later, cells were treated with IFN-β (500pM) or IL-12 (20ng/ml) for 15 min. The level of tyrosine-phosphorylated TYK2 and STAT1 was analyzed with phospho-specific Abs. The membrane was reprobed for TYK2 levels. A nonspecific band shown as loading control. (F) 293T cells were transfected with the pRc-CMV empty vector (EV), TYK2-ΔE8 or the triple mutant TYK2-K930R/Y1044F/Y1045F (DN) possessing dominant-negative activity [19]. Twenty-four hrs later, cells were treated with IFN-α for 15 min. Phosphorylation of STAT1, STAT2 and TYK2 were analyzed with phospho-specific Abs. The membrane strips were reprobed for TYK2 and STAT1 contents. Of note, neither TYK2-ΔE8 nor DN can be inducibly phosphorylated, hence the phospho-TYK2 band corresponds to endogenous TYK2.

To corroborate this finding, we assessed by *in vitro* pull-down assay the ability of the N-ter portions (aa 1-591, FERM and SH2) to interact *in vitro* with the cytoplasmic region of IFNAR1 [22]. N-ter proteins (WT, ΔE8) were incubated with purified GST-IFNAR1cyt and the material retained on beads was analyzed by immunoblot with anti-TYK2 and anti-GST Abs. As shown in Fig 4D, N-ter ΔE8 was not retained on GST-IFNAR1cyt. Overall, the above data indicate that exon 8, which encodes a segment of TYK2 FERM, is essential for the binding of TYK2 to IFNAR1. IL-12 signaling was also assessed. For this, TYK2-WT and TYK2-ΔE8 were transiently transfected in TYK2-null cells expressing the two chains of the IL-12 receptor. As opposed to TYK2-WT, TYK2-ΔE8 failed to rescue IL-12 signaling (Fig 4E). Further, we excluded the possibility that TYK2-ΔE8 abrogates cytokine signaling by acting in a dominant-negative manner (Fig 4F). We thus concluded that TYK2-ΔE8, although well expressed and catalytically competent, is unable to bind cognate cytokine receptors and therefore cannot relay JAK/STAT signaling.

Altogether, the above results demonstrate that the 66 aa segment encoded by exon 8 is critical for cytokine receptor binding. Interestingly, in the solved structure of the FERM-SH2 domain of TYK2 complexed with IFNAR1 (Fig S1B), this segment does not directly contact the receptor [23, 24]. This segment is likely exposed at the surface of the F3 lobe of the FERM and contains a disordered loop (β3-β4 loop), unusually long among the JAK proteins (Fig 1C), which may provide critical contact points for cognate receptors [25].

### Rs2304256 promotes exon 8 inclusion in monocytes

Next, we decided to investigate the impact of the two disease-associated variants on exon 8 by performing a *TYK2 cis*-eQTL analysis at multiple resolution levels. While analysis at whole-gene expression level may mask variation of expression of specific exons, eQTL analysis using RNA-seq data allows quantification of individual splicing events at transcript-, exon-, junction- and intron-levels. We thus used available RNA-seq data from primary CD14^+^ monocytes of 200 genotyped individuals [26]. Interestingly, we found that rs2304256 minor allele A correlated with lower levels of the annotated *TYK2-204* transcript lacking exon 8 (Fig 5A). Consistently, the minor rs2304256 allele also correlated with lower levels of exon 7-exon 9 and exon 7-exon 10 junctions, but higher level of exon 7-exon 8 and exon 8-exon 9 junctions, increased activities of the donor and acceptor sites flanking exon 8, and lower levels of intron 7 and intron 9 (Fig 5, B-D). No correlation with total *TYK2* level was detected (Fig 5E). All of the above most likely reflect the impact of rs2304256 on splicing events resulting on the inclusion of exon 8 and exon 9. Rs12720270 as well correlated to lower expression of *TYK2-204* (P < 0.005), but the effect disappeared when accounting for rs2304256 (P = 0.55) (data not shown). Hence, we could not detect an independent effect of rs12720270 on *TYK2-204* in monocytes.

**Fig 5.**
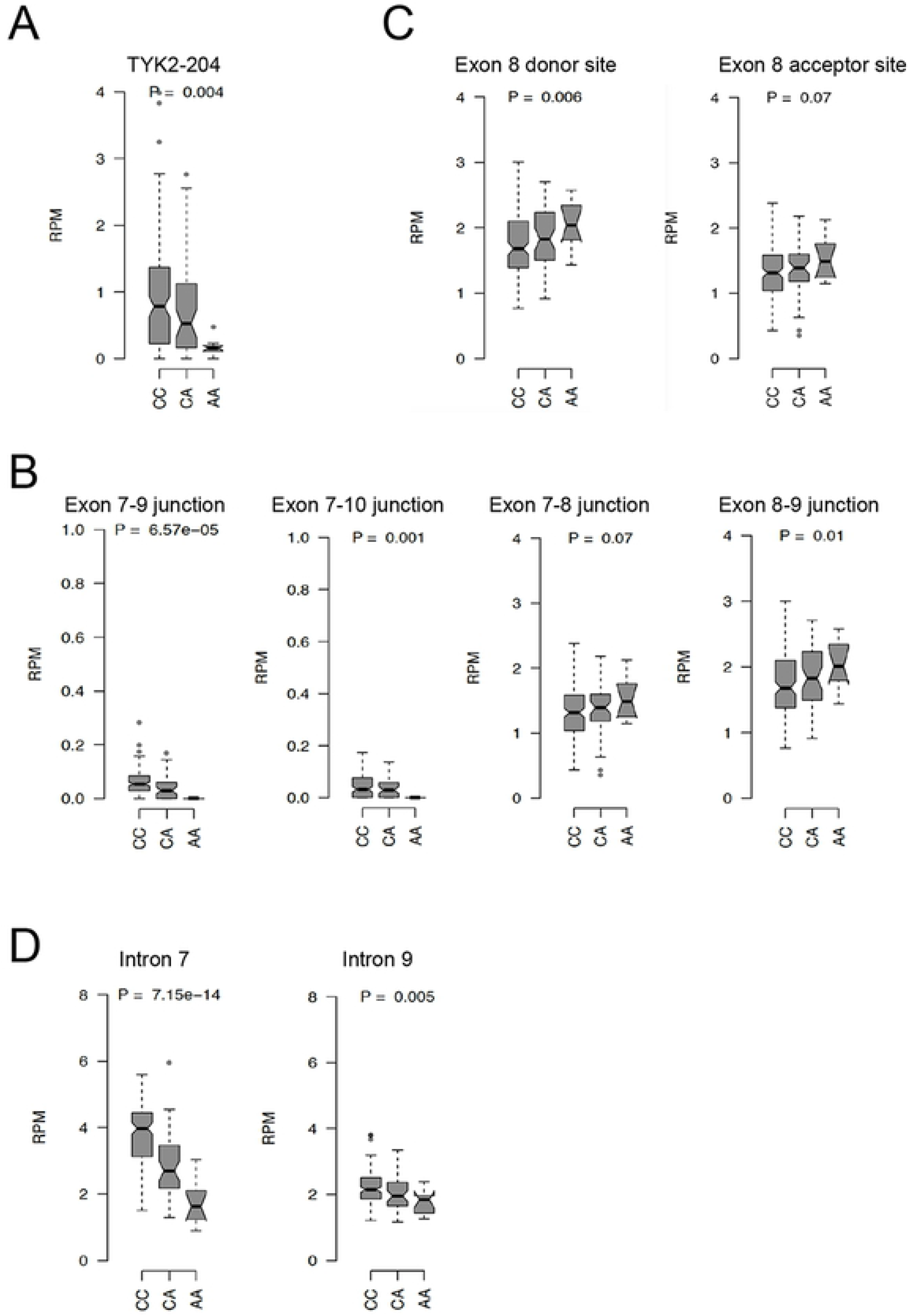
Rs2304256 eQTL analysis on *TYK2* expression in monocytes. Direction of the effect of *cis*-eQTL rs2304256 on *TYK2* in primary CD14+ monocytes from healthy donors (CC n: 143, CA n: 47, AA n: 10). Analysis was done at the level of: (A) transcript, (B) exon-exon junction, (C) splice sites and (D) intron.

### Rs2304256 is associated with increased TYK2

Having shown above that rs2304256 and rs12720270 favour exon 8 inclusion, we wondered if these variants contribute to a higher level of global *TYK2*. Interestingly, in the Genotype-Tissue Expression (GTEx) database, rs2304256 is associated with a modest increase of *TYK2* in several tissues (S1 Table), including whole blood (Fig 6A) (https://gtexportal.org/). We therefore performed an eQTL analysis using *TYK2* NanoString data obtained on whole blood of 1000 healthy donors of European ancestry (Milieu Intérieur cohort) [27, 28]. In non-stimulated blood, rs2304256 showed a tendency toward higher *TYK2*. The effect was modest but significant (P = 0.0048) (Fig 6B).

**Fig 6.**
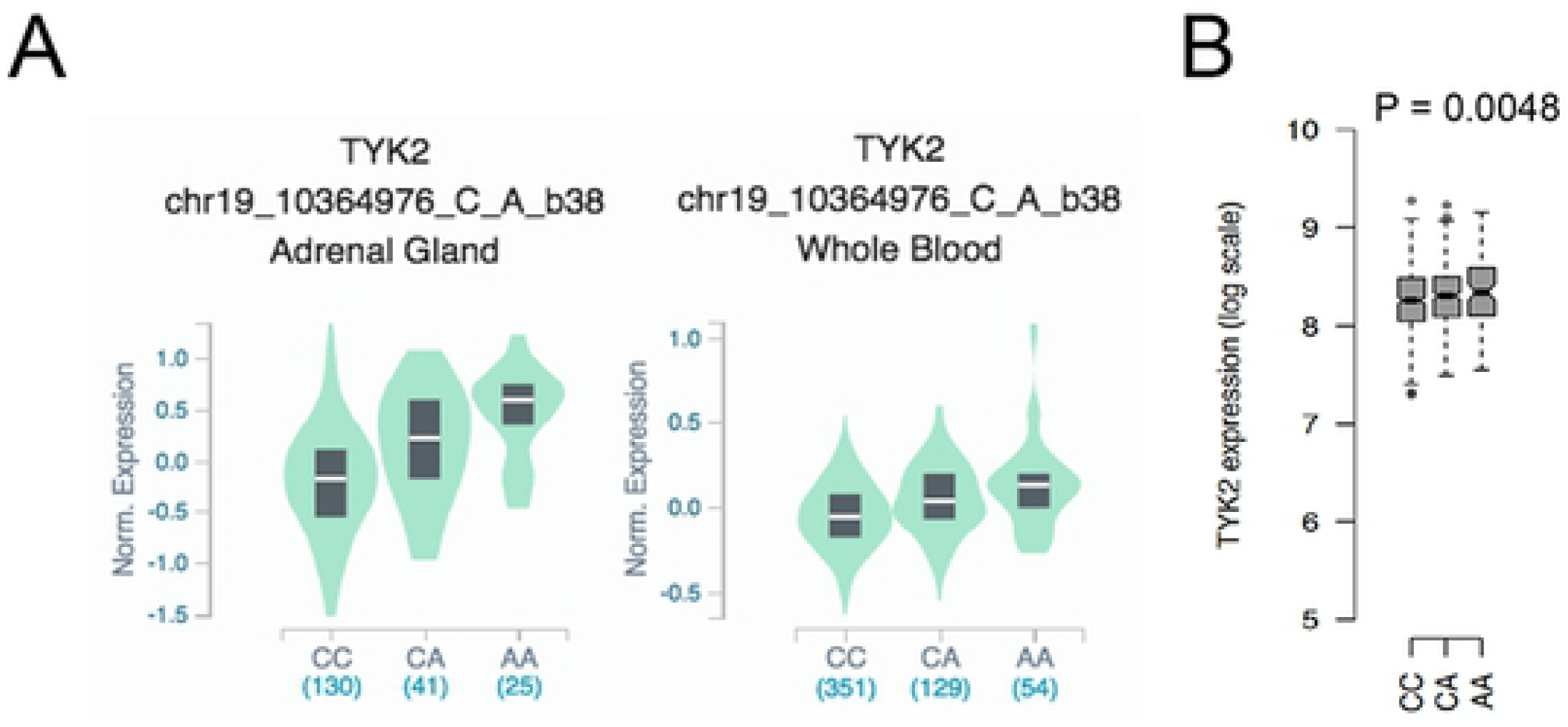
Rs2304256 eQTL analysis on *TYK2* expression in tissues. (A) Direction of effect of *cis*-eQTL of rs2304256 on *TYK2* expression at gene level in adrenal gland and whole blood from GTEx database (https://gtexportal.org/home/snp/rs2304256). (B) Boxplots showing the direction of the effect of *cis*-eQTL of rs2304256 (CC n: 508, CA n: 398, AA n: 81) on *TYK2* expression measured by NanoString in whole blood of 1000 genotyped healthy donors of European ancestry (Milieu Intérieur cohort).

## Discussion

In the last decade, hundreds of loci have been genetically associated to several chronic inflammatory and auto-immune diseases. The complexity of these diseases, our limited knowledge of pathogenic processes and the diverse genetic predisposition still make it difficult to assign causality to single variants in human populations. Several coding variants in the *TYK2* gene have been identified largely through GWAS of populations of European ancestry, and their biological impact and relevance in disease are ill defined. Experimental validation may help to prioritize some of these variants on the basis of their predicted effects at both transcript and protein levels.

In the present study we have analyzed the impact of two common *TYK2* variants that are in strong LD, rs2304256 (V362F) in exon 8 and rs12720270 in intron 7. We found that the amino acid substitution caused by rs2304256 does not alter TYK2 function. *In silico* analyses led us to investigate a possible effect of the two variants on splicing. Analysis of *TYK2* transcripts in genotyped EBV-B cell lines and transcripts expressed from minigene splicing reporters clearly demonstrated the impact of both variants on exon 8, with the minor alleles promoting inclusion. This was validated for rs2304256 in 293T-derived clones that were CRISPR/Cas9-edited at that site. The impact of rs2304256 on exon 8 inclusion was also evident in eQTL analyses of RNA-seq data of primary monocytes. Interestingly, a similar conclusion was reached in the large study by Odhams and collaborators aimed at assessing the causality of SLE-associated *cis*-eQTL variants [29]. These authors performed an eQTL analysis at multiple resolution levels on RNA-seq data from lymphoblastoid cell lines (Geuvadis cohort) and classified rs2304256 as a *cis*-eQTL at the level of exon 8.

Many studies have reported on genetic variants that modulate pre-mRNA splicing and in turn influence phenotype and disease risk [30–32]. These studies can have broad implications as they can uncover pathogenic mechanisms and also inform on therapeutic choice in disease treatment [33–36]. Pre-mRNA splicing is a highly regulated process where each exon is under the combinatorial control of the flanking splice sites and multiple splicing regulatory elements. These elements can be located in exonic and intronic sequences and are not easily identifiable by sequence inspection. They often act as binding platforms for splicing regulators, such as SR proteins and hnRNPs, and can promote exon inclusion (enhancer) or cause exon skipping (silencer). Our results and close sequence analysis suggest that rs2304256 (G > T) generates an ESE motif (CA**<u>T</u>**<u>TCGGC</u>), possibly recognized by SRp55 (SRSF6). The nucleotide variation may at the same time disrupt an ESE recognized by SF2/ASF (<u>CA**G**TCGG</u>C). Interestingly, among the 13 human immune cell types analyzed in the DICE project (https://dice-database.org) [37], SRp55 was found to be particularly abundant in naïve B cells.

The genotyped cell lines and primary cells we have analyzed (carriers of major alleles) exhibit a very low level of the alternate transcript *ΔE8* relative to full-length *TYK2* (Figs 3 and 5). The low-abundance *ΔE8* transcript may result from noisy splicing [38, 39] and may have no biological implication. However, we do not favour this hypothesis, since the alternate *ΔE8* transcript is protein-coding, not subject to nonsense-mediated decay and potentially functional. Alternative splicing is most often cell-specific, tissue-specific or stimulus-induced and the possibility exists that *TYK2 ΔE8* is more abundant and functionally relevant in cells possessing a proper repertoire of splicing regulatory factors. As for the protein, we found that TYK2-ΔE8 has tyrosine kinase activity but cannot bind cytokine receptors, which points to a function unrelated to canonical cytokine signaling. In this regard, we previously reported that overexpressed TYK2 can traffic to the nuclear compartment [21], where it may perform additional roles as proposed for JAK1 and JAK2 [40]. Moreover, TYK2 has been implicated in mitochondrial respiration and differentiation of brown adipose tissue [41, 42]. A non-mutually exclusive possibility is that rs2304256 and rs12720270, by promoting exon 8 inclusion, cause an increase in global (functional) *TYK2* level. Such a ‘dosage’ effect is suggested by GTEx data showing that these polymorphisms correlate with increased *TYK2* level in several tissues but with distinct tissue specificity (S1 Table). (https://gtexportal.org/home/snp/rs2304256) For rs2304256 the best effect size (0.42) is seen in adrenal gland (Fig 6A). Intriguingly, in culture of primary adrenal gland cells, IFN-β was shown to inhibit cortisol production [43]. Hence, the level of TYK2 in adrenal gland may impact on the immune response by modulating cortisol production [44, 45]. Rs2304256, but not rs12720270, correlates with a slightly higher *TYK2* in whole blood (Fig 6B). No such correlation was found in monocytes (data not shown) and lymphoblastoid cells [29], indicating that the impact of rs2304256 on global *TYK2* expression is likely to be cell context-specific.

TYK2 is one of numerous proteins influencing individual susceptibility to chronic diseases with aberrant innate and adaptive immune responses. Since TYK2 transmits signals of a broad range of immuno-regulatory cytokines, natural TYK2 variants may impact a single pathway or a combination of pathways. To date, low-frequency (MAF <5%) coding TYK2 variants associated to protection to AIDs have been shown to relay weaker or no signaling (hypomorphic variants), depending on the cytokine and the cell context [5–7, 9, 10]. Variants increasing TYK2 dosage are expected to lower the threshold of responsiveness - *via* effects on both receptor levels and signaling - thus making cells more sensitive to low cytokine doses. The ultimate impact (protective or damaging) is difficult to predict, as it will depend on disease onset and triggering factors as well as on the cytokines involved in the pathogenic process. TYK2 mediates IL-12 and IL-23 signaling, which can play a pathogenic role in autoimmune and auto-inflammatory conditions. TYK2 also mediates signaling of the anti-inflammatory IL-10, whose protective role is well documented in patients with early-onset IBD and deficiency of IL-10R1 or IL-10R2 [46, 47]. Higher TYK2 dosage may also enhance responsiveness to type I IFN, which exert a broad spectrum of functions and can be pathogenic in many AID. Hence, IFN therapy can induce or exacerbate some AID, such as SLE and T1D [48]. Yet, in the case of fulminant T1D caused by viral infection, high TYK2 in pancreatic β cells may be protective by mediating a robust IFN-mediated antiviral response [49].

In European population studies, rs2304256 was reported to be protective in SLE and other AIDs (Table 1). In subsequent studies, haplotype analysis showed that the rs2304256 association in SLE and RA is likely driven by imperfect LD to the independent causal variants rs34536443 (P1104A), rs12720356 (I684S) or rs35018800 (A928V) [50]. A similar conclusion was reached in a study of systemic sclerosis patients and in a meta-analysis using exome arrays to identify psoriasis-associated rare variants [51, 52]. Recently, a fine mapping analysis of causal variants for RA and IBD identified rs34536443 and rs12720356, but not rs2304256 [53]. Combined with our data, these findings raise the possibility that rs2304256, by acting as a common *cis*-regulatory variant modulating exon 8 inclusion and/or TYK2 dosage, modifies the expressivity of the less common rs34536443 (P1104A) and rs12720356 (I684S) disease variants [54].

Interestingly, in Asian populations rs12720356 (I684S) and rs34536443 (P1104A) are absent or extremely rare, while rs2304256 and rs12720270 are more frequent than in other populations (MAF about 47% and in almost complete LD (r^2^=0.9) (see Table 1). Hence, one would predict that in Asian populations the impact of rs2304256 and rs12720270 on disease susceptibility will not be masked by rs34536443 or rs12720356. In a trans-ancestral GWA meta-analysis, the direction of the effect of rs2304256 on SLE appears opposite in Europeans and Hong Kong Chinese [14]. Yet, results on the association between rs2304256 and SLE in Chinese populations are not consistent [55, 56]. A candidate gene association study on Crohn’s disease in Japanese populations showed that the rs2304256 A allele was significantly more frequent in healthy controls (34.5%) than patients (23.3%) [57]. In two Japanese case/control studies, rs2304256 AA homozygosity was found to be more frequent in SLE patients (17.3%) than controls (13.1%), and in T1D patients (15.2%) than controls (9%) [58, 59]. Additional detailed studies will be necessary to assess the impact of the common rs2304256 variant - particularly in homozygosity - in AIDs in East Asians.

## Materials and methods

### Plasmid constructs

TYK2 WT has been described as pRc-TYK2 [19]. TYK2-P1104A, and TYK2-I684S in pRc-CMV and pIRES vectors respectively, were generated by standard PCR. TYK2-V362, A928V and TYK2-ΔE8 were generated by site-directed mutagenesis using QuickChange XL site-directed mutagenesis kit (Agilent Technologies) in pRc-TYK2 or pQE-His-N [22]. All new plasmids were verified by sequencing. All expression constructs, except TYK2-I684S and pQE-His-N, have a C-terminal vesicular stomatitis virus glycoprotein (VSV-G) epitope tag.

### Cells and transfection

EBV-transformed B cell lines were obtained from Coriell Cell Repositories (Camden, NJ) and from CRB-REFGENSEP (Centre de ressources biologiques du réseau français d’études génétiques sur la sclérose en plaques). Genotyping rs12720270, rs2304256, rs12720356, and rs34536443 confirmed the specific polymorphism in these lines. Cells were cultured in RPMI 1640 and 10% heat-inactivated FCS. The 11,1 (TYK2-deficient fibrosarcoma) and 293T cells were cultured in DMEM and 10% heat-inactivated FCS. Transfections were performed with FuGENE HD (Promega). The 11,1 cells were transfected with pRc-CMV-based plasmids for stable expression of Tyk2 WT or mutants, and clones were selected in 400 µg/ml G418. IFN-α2 was a gift from D. Gewert (Wellcome Research Laboratories).

### Western blot analysis and antibodies

Cells were lysed in modified RIPA buffer (50 mM Tris-HCl pH 8, 200 mM NaCl, 1% NP40, 0.5% DOC, 0.05% SDS, 2mM EDTA) with 1 mM Na_3_VO_4_ and a cocktail of antiproteases (Roche). A total of 30 μg of proteins was separated by SDS-PAGE and analyzed by western blot. Membranes were cut horizontally according to molecular size markers, and stripes were incubated with different Abs. Immunoblots were analyzed by ECL with the ECL Western blotting Reagent (Pierce) or the more sensitive Western Lightning Chemiluminescence Reagent Plus (PerkinElmer) and bands were quantified with Fuji LAS-4000. For reprobing, blots were stripped in 0.2 M glycine (pH 2.5) for 30 min at room temperature. The following Abs were used: TYK2 mAb T10-2 and anti-GST (Hybridolab, Institut Pasteur); anti-IFNAR1 (64G12 mAbs) [60]; anti-STAT2-phospho-Y689 (R&D); Abs to STAT1, STAT2, STAT3, and STAT1-P-Y701, STAT3-P-Y705, and TYK2–P-YY1054/55 (Cell Signaling Technology, Beverly, MA).

### Minigene assay

TYK2 minigene constructs were made in the plasmid vector pI-12 (Addgene). A ∼1.2 kb TYK2 genomic region comprising rs12720270 and rs2304256, spanning from exon 7 to 9, was amplified from genomic DNA of the EBV-B cell line GM52173 using the forward primer 5’-GCCGTCTAGACTTCAAGGACTGCATCCCG-3’ and the reverse primer 5’-GCCGATCGATAGCAGGGGTCCGTGGATC-3’. The PCR product was subcloned into the XbaI and ClaI sites of the pI-12 vector. The different allelic combinations were introduced by site-directed mutagenesis. RNA was isolated from transfected 293T cells using the RNeasy Mini Kit (Qiagen) and cDNA was synthesized using High-Capacity cDNA Reverse Transcription Kit (appliedbiosystems). Minigene splicing was analyzed by PCR amplification of cDNA using T7 and SP6 primers specific to the minigene. All resulting amplification products were sequenced.

### RT-PCR and sequencing of *TYK2* transcripts

To assess the impact of rs2304256 and rs12720270 on splicing of *TYK2* exon 8, we performed RT-PCR and sequenced *TYK2* transcripts that were amplified. Total cellular RNA extraction and reverse transcription were performed using the method described above. We used the forward primer 5’-GCCGTCTAGACTTCAAGGACTGCATCCCG-3’ and reverse primer 5’-GCCGATCGATAGCAGGGGTCCGTGGATC-3’ to amplify transcripts containing exons 7, 8 and 9, and the forward primer 5′-GAGTCATCGCTGACAACTGAGGAAGTCTGCATC-3′ and reverse primer 5′-GCACAGGTAGTGGCTGGAG -3′ to amplify all transcripts that contained from exon 3 to exon 9. All PCR products were separated and purified from agarose gel before being sent for sequencing.

### CRISPR-Cas9-modified cell lines

293T cells were modified using the CRISPR-Cas9 system as in [61]. We used the plasmid-based delivery method. Briefly, to introduce the rs2304256 minor allele, the oligo pair encoding the guide sequence (forward 5′-CACCGCCAAGGCTCACAAGGCAGT-3′, reverse 5′-AAACACTGCCTTGTGAGCCTTGGC -3′) were annealed and ligated into the vector pSpCas9(BB)-2A-Puro (PX459) for co-expression with Cas9, a gift from Feng Zhang (Addgene plasmid 62988). The homology-directed repair (HDR) template (5’cacttgctgggtgttcagGGTTCTAGTGGCAGCAGTGGCAGGAACCCCCAAGCCAGCCTGTTTGGGAAGAAGGCCAAGGCTCACAAGGCATTCAGCCAGCCGGCAGACAGGCCGCGGGAGCCACTGTGGGCCTACTTCTGTGACTTCCGGGACATCACCCACGTGGTGCTGAAAGAGCACT-3’) was co-transfected in 293T cells with the above sgRNA expression plasmid using Fugene HD. Cells were kept in puromycin (0.9 μg/ml) for 4 days. Individual clones were genotyped for rs2304256 by Sanger sequencing. *TYK2*-knockout cell lines were generated by transfecting 293T cells with sgRNA expression plasmid without HDR template.

### *In vitro* kinase assay

Cells were lysed in 50 mM Tris (pH 6.8), 0.5% Nonidet P-40, 200 mM NaCl, 10% glycerol, 1 mM EDTA, 1 mM sodium vanadate, 1 mM sodium fluoride,10 mM PMSF, cocktail of antiproteases (Roche Applied Science). TYK2 and JAK1 were immunoprecipitated from 2 mg lysate using affinity-purified anti-VSV-G polyclonal Abs (a gift from M. Arpin, Institut Curie). Immunocomplexes were washed three times in buffer 1 (50 mM Tris [pH 6.8], 400 mM NaCl, 0.5% Triton X-100, and 1 mM EDTA), once in buffer 2 (50 mM Tris [pH 6.8] and 200 mM NaCl), and once in kinase buffer (50 mM HEPES [pH 7.6] and 10 mM MgCl_2_). The kinase reaction was carried out in 50 mM HEPES [pH 7.6], 10 mM MgCl_2_ and with or without 30 μM ATP at 30°C for 5 min in a total volume of 30 µl. The reaction was terminated by boiling in Laemmli buffer. Half of the sample was loaded for SDS-PAGE, transferred to a nitrocellulose membrane, and phosphorylated products were analyzed by western blotting with activation loop phospho-specific Abs. After stripping, membranes were reblotted with anti-TYK2 mAb, revealed using ECL detection reagents (Western Lightning, PerkinElmer).

### Protein purification and *in vitro* pull-down assay

Histidine-tagged proteins were expressed in bacteria, purified on Ni-NTA agarose beads according to the manufacturer’s protocol (Qiagen), eluted, and dialyzed against 20 mM Tris-HCl, pH 7.5, 100 mM NaCl, 10% glycerol, and 2mM EDTA, 1mM dithiothreitol (DTT). Proteins were concentrated by Vivaspin concentrator (Vivascience) and stored at 80 °C. GST fusion proteins were affinity-purified on glutathione-Sepharose (GE Healthcare). For *in vitro* pull-down assays, same quantity of His-tagged purified recombinant proteins were incubated with glutathione-Sepharose containing about 2μg of bound GST fusion protein in 100 μl of binding buffer (0.1% Nonidet P-40, 10% glycerol, 50 mM NaCl, 50 mM Tris-HCl, pH 8, and 1 mM DTT) with 0.5% bovine serum albumin and protease inhibitors for 60 min at 4 °C. Beads were pelleted and washed three times in binding buffer. Bound proteins were eluted and boiled in 20 μl of Laemmli buffer, separated by SDS-PAGE, and analyzed by immunoblotting with the appropriate antibody.

### Quantification and statistical analysis

RNA-Sequencing (RNA-Seq) data on *TYK2* expression in primary monocytes derived from 200 genotyped healthy male individuals of self-reported African and European ancestry are from the EvoImmunoPop project [26], and were analyzed as described in [38, 39] at five levels: gene-, transcript-, exon-, junction-, and intron-level. Expression data are corrected for sequencing depth and gene/transcript/exon/intron length (RPKM for transcripts, exons, introns and RPM for exon-exon junctions and splicing sites). Briefly, transcripts were quantified with Cuffdiff [62], based on Ensembl v70 annotations. For exon and intron quantification, all exons were split into non-overlapping exonic parts [63], and a pseudo-transcript was build containing the union of all exons from the gene. Genic regions located between exons from the pseudo-transcript were then used as introns. Quantification of gene expression from exonic parts and introns was done using HT-Seq [64]. For the quantification of splice junction, we used the the filter_cs script from leafcutter package [65] to extract all spliced reads with an overhang of at least 6 nucleotides into each exon and count reads across each exon-exon junction. EQTL analyses were performed using Matrix EQTL [66], including for population of origin as a covariate. *TYK2* mRNA levels were measured in whole blood of healthy donors from the Milieu Intérieur cohort by the NanoString hybridization-based multiplex assay [28]. The Milieu Intérieur cohort is composed of 500 men and 500 women from 20 to 69 years of age [67, 68]. The NanoString *TYK2* probe is complementary to the exon 3/exon 4 junction: GGGCCTGAGCATCGAAGAGGGCAAAGAGATTGAAGCAAGGAGGAGTGATACCAACTTTATGTGCAATGTGGATGCAGACTTCCTCAGCTGTCAGCGATGA. eQTL analyses were performed with the linear mixed model implemented in GenABEL R package [69].

## Supporting information

**S1 Table. Summary of single-tissue eQTL for rs2304256 and rs12720270 on TYK2 transcript expression across 48 tissues**

**S1 Fig. Skipping of exon 8 maintains the correct reading frame of TYK2**. Partial exon 7 and exon 9 sequences are boxed. The central exon 8 encodes the 66 aa segment. Red arrowhead points to Val362.

**S2 Fig. The exon 8-encoded segment within the TYK2 FERM**. Taken and adapted from Supplem. Fig. S3 in (Wallweber HJ et al, Nat. Struct. Mol. Biol. 2014. 21:443). Shown is the secondary structure of the TYK2 FERM-SH2 domain in complex with a short peptide of the IFNAR1 cytoplasmic portion (yellow). SH2 domain in light blue. The FERM comprises three lobes (F1, F2 and F3). Helices displayed as cylinders, strands displayed as block arrows, and loops displayed as lines. Arrows point to the start (5’) and the end (3’) of the exon 8-encoded segment, spanning from the β3-β4 loop to the β7 strand of the F3 lobe. Valine 362 is located in the unstructured β3-β4 loop and is tentatively indicated by an arrow.

## Acknowledgments

We wish to thank Barbara Piasecka for help in initial analyses of *TYK2* variants in the Milieu Intérieur cohort.

## Author contributions

ZL designed and performed experiments. ZL, FM and SP analyzed the data. MR and EP performed the eQTL analyses. ZL, SP, MR wrote and edited the manuscript. FM advised and revised the manuscript. SP supervised the work.

## Notes

**Funding**: Research in the Unit of Cytokine Signaling funded by the Institut Pasteur, the Fondation pour la Recherche Médicale (Equipe FRM DEQ20170336741) and the Institut National de la Santé et de la Recherche Médicale (INSERM). ZL is supported by the Centre National de la Recherche Scientifique (CNRS). Research in the Unit of Human Evolutionary Genetics funded by the Institut Pasteur, the French Government’s Investissement d’Avenir program, Laboratoires d’Excellence “Integrative Biology of Emerging Infectious Diseases” (ANR-10-LABX-62-IBEID), “Milieu Intérieur” (ANR-10-LABX-69-01) and the Fondation pour la Recherche Médicale (Equipe FRM DEQ20180339214).

